# Faster responses of photosynthesis to light transitions increase biomass and grain yield in transgenic *Sorghum bicolor* overexpressing Rieske FeS

**DOI:** 10.1101/2022.07.25.501469

**Authors:** Maria Ermakova, Russell Woodford, Zachary Taylor, Robert T. Furbank, Srinivas Belide, Susanne von Caemmerer

**Author notes:** Corresponding author: Maria Ermakova,. Susanne von Caemmerer.

## Abstract

Sorghum is one of the most important crops providing food and feed in many of the world’s harsher environments. Sorghum utilises the C_4_ pathway of photosynthesis in which a biochemical carbon concentrating mechanism results in high CO_2_ assimilation rates. Overexpressing the Rieske subunit of the Cytochrome *b*_6_*f* complex was previously shown to increase the rate of photosynthetic electron transport and stimulate CO_2_ assimilation in the model C_4_ plant *Setaria viridis*. To test whether productivity of C_4_ crops could be improved by Rieske overexpression, we created transgenic *Sorghum bicolor* plants with increased Rieske content. The transgenic plants showed no marked changes in abundance of other photosynthetic proteins or chlorophyll content. Increases in yield of Photosystem II and CO_2_ assimilation rate as well as faster responses of non-photochemical quenching during transient photosynthetic responses were observed as a result of an elevated *in vivo* Cytochrome *b*_6_*f* activity in plants overexpressing Rieske. The steady-state rates of electron transport and CO_2_ assimilation did not differ between transgenic and control plants, suggesting that Cytochrome *b*_6_*f* is not the only factor limiting electron transport in sorghum at high light and high CO_2_. Nevertheless, more agile responses of photosynthesis to light transitions led to increases in biomass and grain yield in plants overexpressing Rieske. Our results indicate that increasing Rieske content could boost productivity of C_4_ crops by improving the efficiency of light utilisation and conversion to biomass.

## Introduction

C_4_ plants utilise a specialised photosynthetic pathway in which a metabolic C_4_ cycle acts as a biochemical carbon concentrating mechanism (Hatch, 1987). The C_4_ cycle operates between mesophyll and bundle sheath (BS) cells, and Ribulose-1,5-bisphosphate carboxylase/oxygenase (Rubisco), the main enzyme of CO_2_ fixation, is localised to the BS (Kanai and Edwards, 1999). Atmospheric CO_2_ (in the form of HCO_3-_) is first fixed in mesophyll cells by PEP carboxylase (PEPC) into a C_4_ acid (hence the term C_4_ photosynthesis). C_4_ acids diffuse to BS cells where they are decarboxylated to produce pyruvate and CO_2_, providing high CO_2_ partial pressure (*p*CO_2_) around Rubisco (Furbank and Hatch, 1987). Higher carboxylation efficiency of Rubisco in C_4_ plants allows higher radiation use efficiency and increased biomass production compared to C_3_ plants (Long, 1999; Sage and Zhu, 2011). Because of their superior productivity, C_4_ crops are becoming increasingly important for food and bioenergy security. The global production of C_4_ maize (*Zea mays*) often surpasses the two key C_3_ cereals, wheat and rice, and C_4_ miscanthus (*Miscanthus* × *giganteus*) and switchgrass (*Panicum virgatum*) are two of the currently leading dedicated biomass crops. This has created considerable interest in identifying and testing strategies to improve productivity of C_4_ crops (Sales et al., 2021; von Caemmerer and Furbank, 2016).

While C_4_ plants are more productive, running the C_4_ cycle requires additional input of energy. Whilst C_3_ plants need at least two NADPH and three ATP to fix one mol of CO_2_, C_4_ plants need two additional ATP molecules to regenerate PEP from pyruvate in mesophyll cells (Edwards et al., 2001; Hatch, 1987). NADPH and ATP are the products of light reactions of photosynthesis which include electron and proton transport in the thylakoid membranes of chloroplasts. NADPH is produced during linear electron flow as electrons originating from water split by Photosystem II (PSII) are transferred by the chain of cofactors, via Cytochrome *b*_6_*f* complex (Cyt*b*_6_*f*) and Photosystem I (PSI), to NADP^+^. Cyt*b*_6_*f* links oxidation of plastoquinol with the translocation of protons to the lumen, a space enclosed by the thylakoid membrane, by operating the Q-cycle (Malone et al., 2021). The transmembrane proton gradient (ΔpH) established across the thylakoid membrane creates a proton motive force (*pmf*) that drives ATP synthesis via the ATP synthase complex. In addition to linear electron flow, C_4_ plants run active cyclic electron flow to produce additional ATP (Ishikawa et al., 2016; Munekage and Taniguchi, 2016; Nakamura et al., 2013). Cyclic electron flow returns electrons from the reducing side of PSI back to the plastoquinone (PQ) pool to repeat plastoquinol oxidation by Cyt*b*_6_*f* and build up additional *pmf* (Johnson, 2011). Thus, cyclic electron flow results in the net production of ATP but not NADPH. ΔpH controls PSII activity by regulating the energy-dependent and quickly reversible form of non-photochemical quenching (NPQ) (Li et al., 2002). The latter is a common term for diverse reactions that help to reduce excitation energy reaching reaction centers of PSII (Malnoë, 2018). Establishing energy-dependent NPQ requires the PsbS protein that senses luminal pH and modifies the light-harvesting complex II (LHCII) to dissipate a part of absorbed light as heat, as well as the conversion of violaxanthin to zeaxanthin (Johnson et al., 2009; Li et al., 2004).

The vast majority of agriculturally important C_4_ crops, like maize, sorghum (*Sorghum bicolor*), sugarcane, miscanthus and several millets (*e*.*g*., *Setaria italica*), belong to the NADP-ME subtype of C_4_ photosynthesis which employs NADP^+^-dependent malic enzyme to decarboxylate the C_4_ acid malate in BS chloroplasts (Furbank, 2011). Because of the drastic differences in biochemistry of mesophyll and BS cells in NADP-ME plants, electron transport chains of the two cell types are also largely different: mesophyll cells predominantly run linear electron flow and BS cells – cyclic electron flow (Ermakova et al., 2021a; Munekage, 2016). To coordinate the production of NADPH and ATP between cells, plants need to tightly regulate the distribution of available light energy (Bellasio and Ermakova, 2021; Bellasio and Lundgren, 2016). Because of their higher ATP requirement, C_4_ plants generally require higher solar radiation and are typically found in tropical regions (Sage et al., 2011). At low irradiance however, slow operation of the C_4_ cycle results in a lower *p*CO_2_ in BS cells decreasing the efficiency of C_4_ photosynthesis (Furbank and Hatch, 1987; Kromdijk et al., 2010). Therefore, increasing radiation use efficiency is one of the primary strategies for increasing assimilation rates and productivity of C_4_ plants.

Constitutive overexpression of the Rieske FeS subunit of Cyt*b*_6_*f* (hereafter Rieske), encoded by the nuclear *petC* gene was shown to increase abundance of the whole complex in both mesophyll and BS cells of a model NADP-ME grass *Setaria viridis* (Ermakova et al., 2019). This resulted in a higher quantum yield of both photosystems and higher CO_2_ assimilation rates at high light and high CO_2_. However, the feasibility of using Rieske overexpression for improving crop productivity required further assessment. Here we test effects of Rieske overexpression on biomass and grain yield of a multipurpose C_4_ crop, sorghum. We show that sorghum plants with increased Rieske abundance use light more efficiently and accumulate more biomass due to faster responses of photosynthesis to light transitions. Our results indicate that increasing Rieske content is a promising strategy for stimulating yield of sorghum and other C_4_ crops.

## Results

Sorghum plants transformed with the construct for Rieske overexpression (see Materials and Methods for details) were selected based on kanamycin resistance and transferred to soil for growth in a glasshouse. Sixteen T_0_ plants were recovered and analysed for insertion number, transgene expression and leaf Rieske abundance. The *ntpII* insertions and transcripts of *Brachipodium dystachion petC* (*BdpetC*) were confirmed in ten T_0_ plants (Fig. 1a and Fig. 1b). Plants 25, 26, 29 and 32 showed relatively higher Rieske abundance per leaf area compared to wild type (WT) and escape plants without the T-DNA insertion (Fig. 1a). The T_1_ progenies of those four plants were grown and analysed for insertion numbers and Rieske abundance. The homozygous T_1_ plants of lines 25 and 26 (4 insertions, Fig. 1c) had higher Rieske leaf content compared to control plants (WT and null segregants). Furthermore, homozygous plants 25-11 and 26-11 showed relatively lower NPQ compared to control and other T_1_ plants when assayed at ambient light, in line with the NPQ phenotype reported in *S. viridis* overexpressing Rieske (Ermakova et al., 2019). The progenies of those two plants, as well as the progeny of the homozygous T_1_ plant 32-14 with increased Rieske abundance (Fig. S1), were used in further experiments, and are hereafter referred to as transgenic lines 25, 26 and 32.

**Fig. 1.**
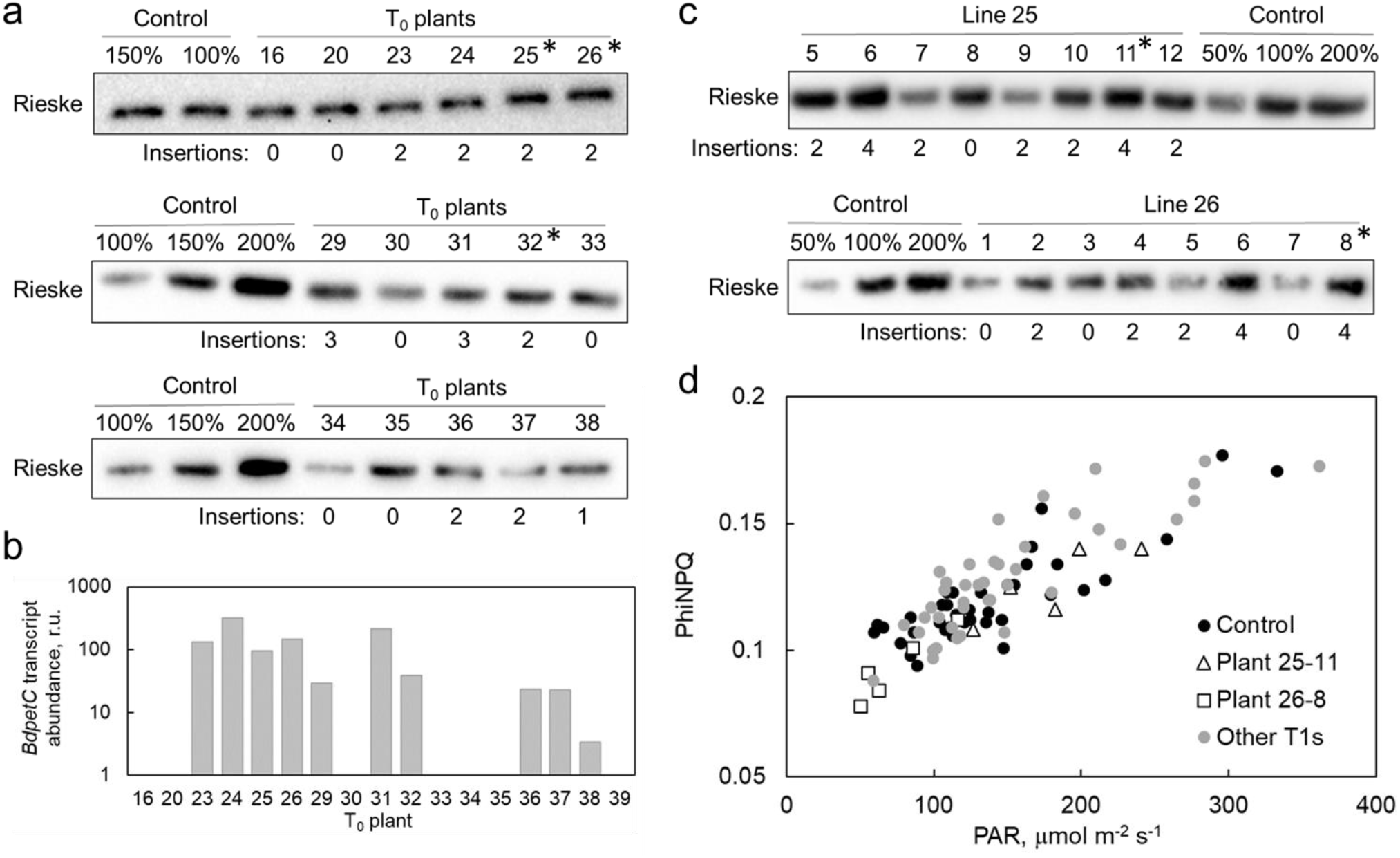
Selection of transgenic sorghum lines overexpressing Rieske. **a**. Immunodetection of Rieske in leaves of T_0_ plants. **b**. Transcript abundance of *B. dystachion petC* (*BdpetC*) in T_0_ sorghum plants. **c**. Immunodetection of Rieske in T_1_ progenies of lines 25 and 26. (**a** and **c**) Samples were loaded on leaf area basis, and the titration series of WT samples was used for relative quantification. Insertion numbers indicate a copy number of *ntpII* obtained by digital PCR. Asterisks indicate the plants which progenies were used in further experiments. **d**. Quantum yield of non-photochemical quenching (PhiNPQ) measured at ambient irradiance in T_1_ progenies of lines 25 and 26. WT and azygous plants were used as control. Each point represents a technical replicate.

T_2_ plants of the three transgenic lines and azygous control plants were grown over summer in a glasshouse with natural light. Abundance of photosynthetic proteins was analysed in leaf extracts loaded on leaf area basis by immunoblotting with specific antibodies (Fig. 2a). Quantification of immunoblots demonstrated a significant, about 40%, increase of Rieske content in all three transgenic lines compared to control plants. Relative abundance of other electron transport components, such as the D1 protein of PSII, AtpB subunit of ATP synthase, PsbS and Lhcb2 subunit of LHC II was largely unaltered in transgenic plants overexpressing Rieske (Fig. 2b) as well as the relative Chl content (Table 1). The content of PEPC and Rubisco large subunit (RbcL) did not differ between transgenic and control plants (Fig. 2b).

**Fig. 2.**
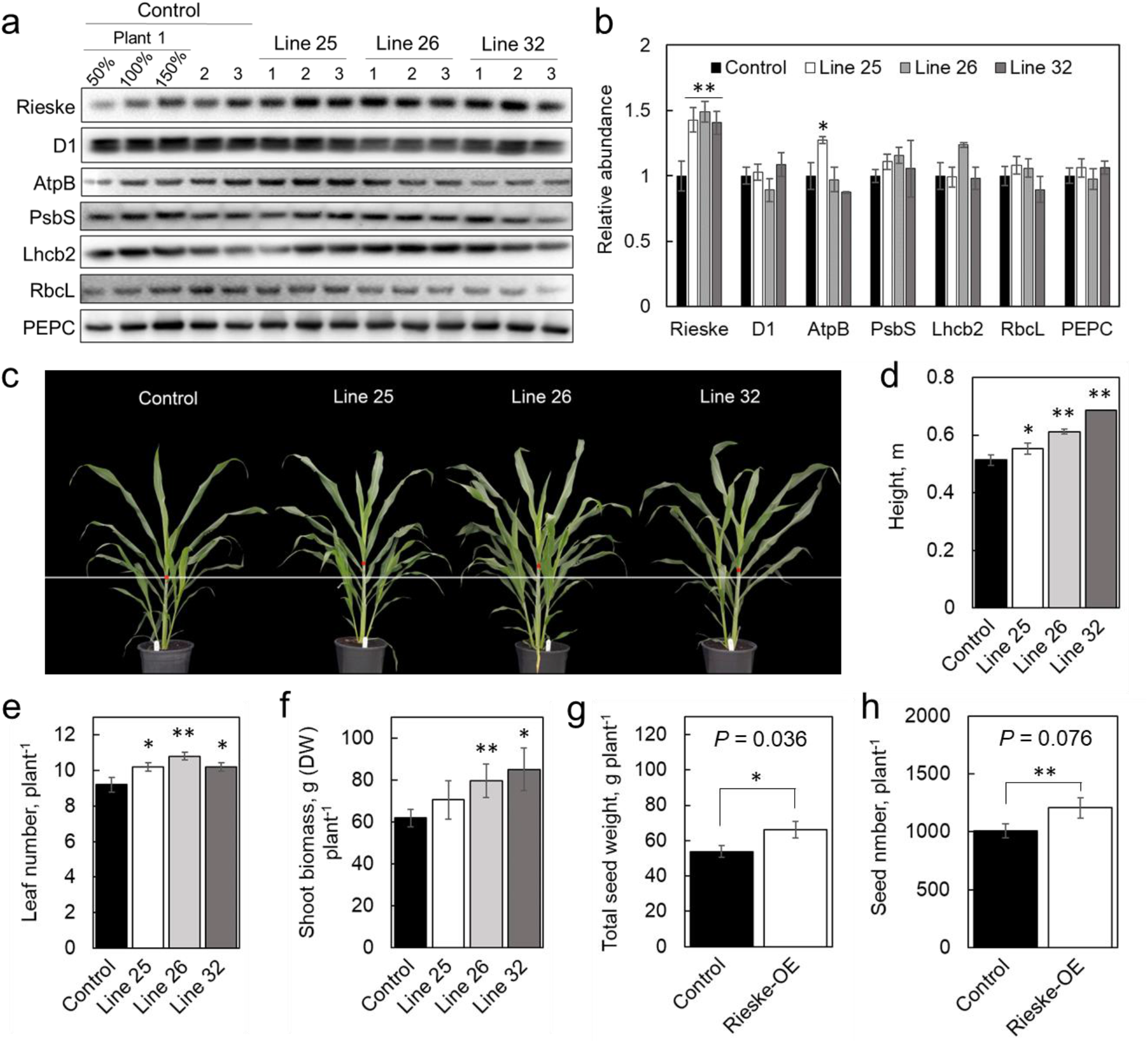
Protein analysis and growth of sorghum lines overexpressing Rieske. **a**. Immunodetection of photosynthetic proteins in leaves of control and transgenic plants: Rieske (Cyt*b*_6_*f*), D1 (PSII core), AtpB (ATP synthase), PsbS (energy-dependent non-photochemical quenching), Lhcb2 (light-harvesting complex II), RbcL (large subunit of Rubisco), PEPC (PEP carboxylase). Samples were loaded on leaf area basis, and the titration series of the Control sample #1 was used for relative quantification. **b**. Quantification of immunoblots relative to control plants. Mean ± SE, *n* = 3 biological replicates. **c**. Phenotype of plants 5 weeks after germination. (**d, e, f**) Height, leaf number and plant biomass, *n* = 5 biological replicates. (**g, h**) Total weight and number of seeds produced per plant, *n* = 18 biological replicates for Control (WT and azygous plants), *n* = 20 for Rieske-OE (lines 25 and 26). Asterisks indicate statistically significant differences between transgenic and control plants based on one-way ANOVA or *t*-test (\*\**P* < 0.05 or \**P* < 0.1).

**Table 1.** Properties of control plants and transgenic sorghum lines overexpressing Rieske. Azygous plants were used as control. Mean ± SE, *n* = 5 biological replicates. Asterisks indicate statistically significant differences between transgenic and control plants (one-way ANOVA and Dunnett’s *post-hoc* test at *P* < 0.05). LMA, leaf mass per area; F_V_/F_M_, the maximum quantum efficiency of PSII.

|  | Control | Line 25 | Line 26 | Line 32 |
| --- | --- | --- | --- | --- |
| Relative Chl | 60.1 ± 1.9 | 57.6 ± 1.9 | 60.9 ± 1.8 | 60.5 ± 1.8 |
| Leaf thickness, mm | 0.63 ± 0.05 | 0.63 ± 0.03 | 0.60 ± 0.03 | 0.63 ± 0.04 |
| LMA, g (dry weight)<br>m<sup>-2</sup> | 119.42 ± 4.78 | 115.73 ± 4.08 | 113.92 ± 4.72 | 122.32 ± 2.65 |
| Tiller number, plant<sup>-1</sup> | 1.4 ± 0.27 | 2.4 ± 0.45 | 2.0 ± 0.36 | 2.0 ± 0.36 |
| $F_V/F_M$ | 0.802 ± 0.002 | 0.802 ± 0.006 | 0.803 ± 0.003 | 0.810 ± 0.001* |

Sorghum plants with increased Rieske content were taller than the control plants at 5 weeks after germination (Fig. 2c and Fig. 2d) and had more tillers (0.036 < *P* > 0.09, Table 1) and leaves (Fig. 2e). Whilst the leaf thickness and leaf dry mass per area did not differ between the genotypes (Table 1), the total aboveground biomass of lines 26 and 32 at harvest was higher compared to control plants (Fig. 2f). In another experiment, when plants were grown in a glasshouse during late summer-autumn, Rieske-OE plants of lines 25 and 26 had larger leaves compared to control plants (azygous and WT, Fig. S2) and produced more seeds by weight and number than control plants (Fig. 2g and Fig. 2h).

Next, we analysed photosynthetic properties of sorghum plants overexpressing Rieske. First, we conducted gas exchange and fluorescence analysis at different *p*CO_2_ and irradiances. No significant differences in CO_2_ assimilation rate or the effective yield of PSII (PhiPSII) were detected between the plants overexpressing Rieske and control plants at constant irradiance of 1500 µmol m^-2^ s^-1^ and different *p*CO_2_ (Fig. 3, left panels). At ambient *p*CO_2_, CO_2_ assimilation rates and stomatal conductance were similar between the genotypes at all irradiances (Fig. 3, right panels). The photochemical and non-photochemical yields of PSI and PSII analysed at different irradiances were largely unaltered in transgenic plants compared to control plants (Fig. 4), except for the yield of PSI (PhiPSI) being higher and the non-photochemical loss of PSI yield due to the acceptor side limitation (PhiNA) being lower in lines 25 and 32 at 95 µmol m^-2^ s^-1^. These results indicated that the steady-state rates of electron transport and CO_2_ assimilation were largely not affected in plants overexpressing Rieske. The maximum quantum efficiency of PSII (F_V_/F_M_), however, was significantly higher in plants of line 32 compared to control plants (Table 1).

**Fig. 3.**
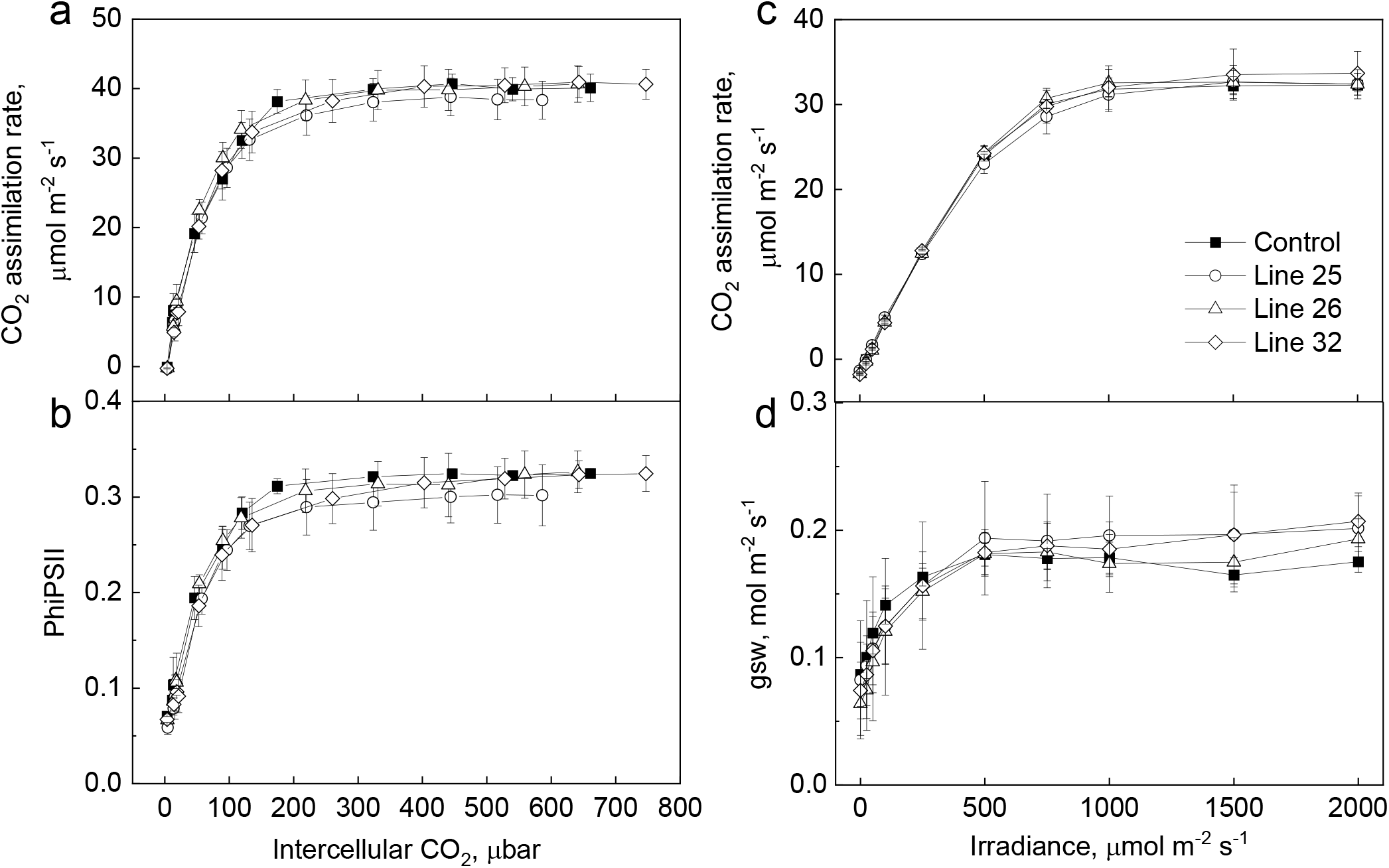
Gas exchange and fluorescence analysis of control and transgenic sorghum plants overexpressing Rieske at different *p*CO_2_ (left panels, measured at 1500 µmol photons m^-2^ s^-1^) or irradiance (right panels, measured at ambient *p*CO_2_). PhiPSII, the effective quantum yield of PSII; gsw, stomatal conductance to water vapor. Azygous plants were used as control. Mean ± SE, *n* = 5 biological replicates. No statistically significant differences were found between transgenic and control plants (one-way ANOVA and Dunnett’s post-hoc test at *P* < 0.05).

**Fig. 4.**
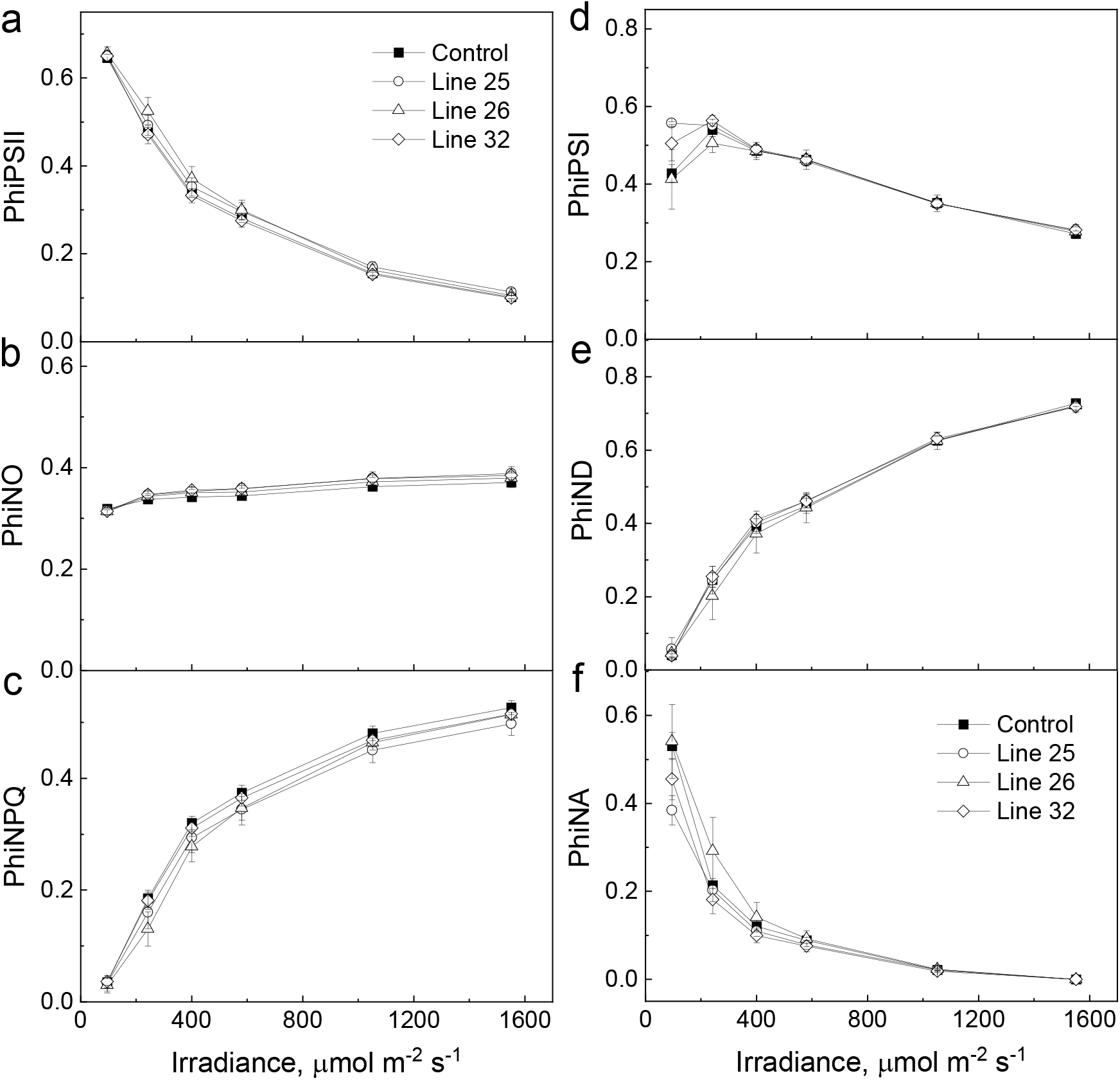
Electron transport parameters estimated at different irradiance from leaves of control and transgenic sorghum plants overexpressing Rieske. PhiPSII, the effective quantum yield of PSII; PhiNPQ, the yield of non-photochemical quenching; PhiNO, the yield of non-regulated non-photochemical reactions within PSII; PhiPSI, the effective quantum yield of PSI; PhiND, the non-photochemical yield of PSI due to the donor side limitation; PhiNA, the non-photochemical yield of PSI due to the acceptor side limitation. Azygous plants were used as control. Mean ± SE, *n* = 5 biological replicates. Asterisks indicate statistically significant difference between line 25 (black) or line 32 (grey) and control plants (one-way ANOVA and Dunnett’s post-hoc test at *P* < 0.05).

Energisation properties of the thylakoid membranes were tested by recording electrochromic shift signal and absorbance changes at 535 nm. By the end of 3-min illumination intervals, *pmf* and proton conductivity of the thylakoid membrane (*g*_H+_), reflecting the speed of *pmf* dissipation and thus ATP synthase activity, did not differ between the plants overexpressing Rieske and control plants at any irradiance (Fig. 5a and Fig. 5b). To gain information about the kinetics of ΔpH build-up during the illumination, we recorded absorbance changes at 535 nm which reflect both zeaxanthin formation and the LHCII modifications induced by PsbS, therefore, providing information on the response of NPQ to ΔpH (Horton et al., 1991; Li et al., 2004). All three transgenic lines overexpressing Rieske established NPQ significantly faster than control plants in the beginning of illumination upon the shift from dark to 1600 µmol m^-2^ s^-1^, indicating a transiently faster build-up of ΔpH due to the increased Cyt*b*_6_*f* activity (Fig. 5c).

**Fig. 5.**
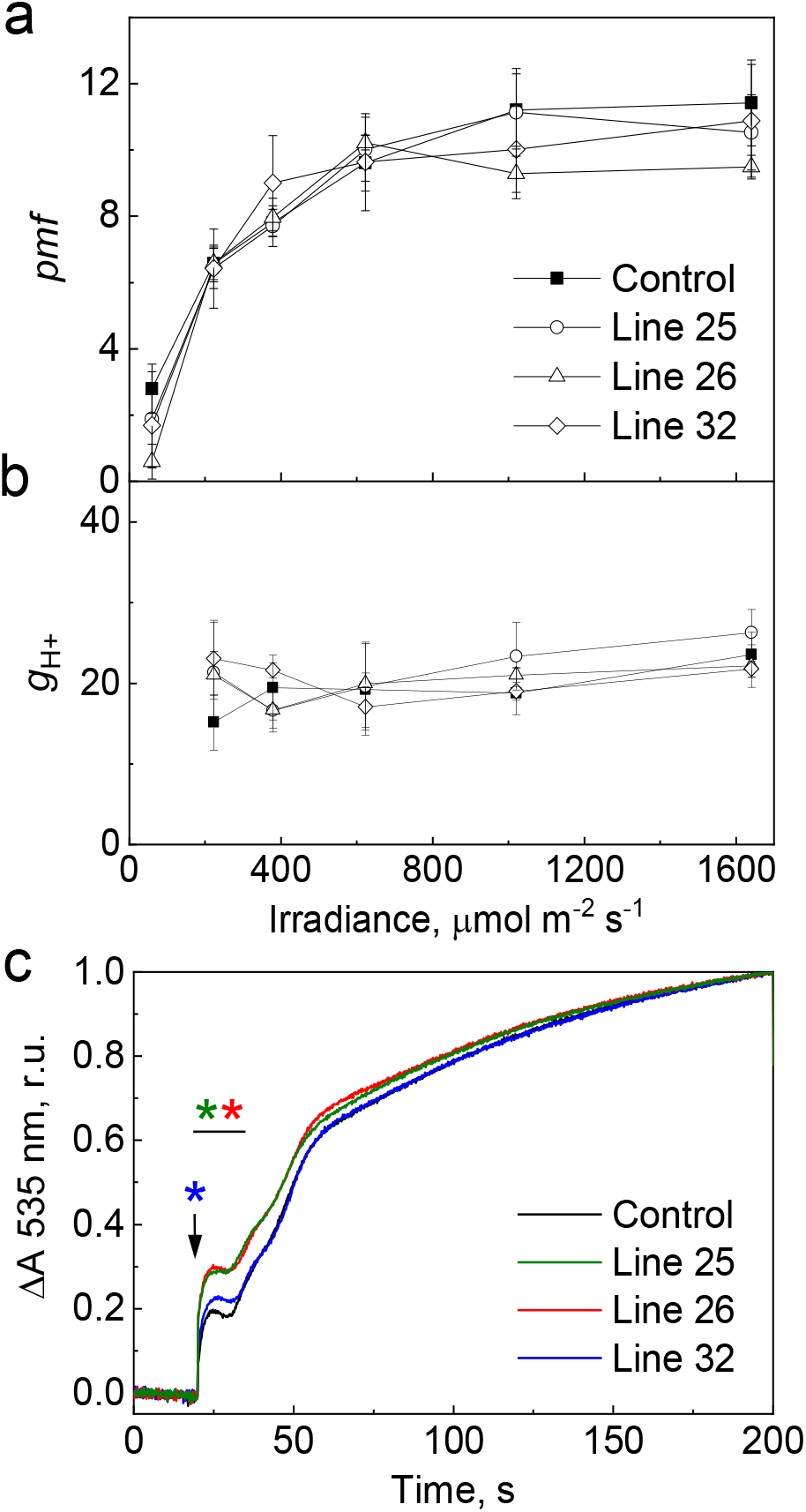
Analysis of the thylakoid membrane energisation in control plants and transgenic sorghum lines overexpressing Rieske. (**a** and **b**) Proton motive force (*pmf*) and proton conductivity of the thylakoid membrane (*g*_H+_) at different irradiance analysed using electrochromic shift signal. Mean ± SE, *n* = 4 biological replicates. **c**. Absorbance changes at 535 nm recorded upon the shift from dark to 1600 µmol m^-2^ s^-1^. The absorbance at the beginning and end of the 3-min illumination period was normalised to 0 and 1, respectively, to facilitate comparison of the kinetics. Averages of 4 biological replicates are presented. Azygous plants were used as control. Asterisks indicate intervals of statistically significant difference between transgenic and control plants (one-way ANOVA and Dunnett’s post-hoc test at *P* < 0.05).

Since transient increase of Cyt*b*_6_*f* activity could be detected upon changes in illumination, we analysed the induction of photosynthesis in overnight-dark-adapted plants during the first 30 min of illumination with actinic light of 1000 µmol m^-2^ s^-1^ (Fig. 6). Because the steady-state CO_2_ assimilation rate, PhiPSII and NPQ did not differ between genotypes (Fig. 3 and Fig. 4), we normalised these parameters to minimum and maximum values to facilitate comparison of the kinetics. During the induction of photosynthesis, CO_2_ assimilation rates increased faster in sorghum plants overexpressing Rieske compared to control plants, and between 18 and 26 min the rates were significantly higher in all three transgenic lines. The induction kinetics of PhiPSII was similar to the kinetics of CO_2_ assimilation rate. Transgenic lines reached the steady-state faster and lines 25 and 26 had significantly increased PhiPSII between 18 and 26 min since the beginning of illumination, compared to control plants (Fig. 6). Interestingly, during the first 9 min of illumination plants overexpressing Rieske built-up NPQ faster than control plants (significant in line 25 and 26) and line 32 then relaxed NPQ significantly faster. Importantly, although the stomatal conductance was significantly higher in all three lines overexpressing Rieske (Fig. 6d), increased CO_2_ assimilation rates were not caused by an increased CO_2_ availability since the ratio of intercellular to ambient CO_2_ partial pressures (*C*_i_/*C*_a_) was unaltered (Fig. 6).

**Fig. 6.**
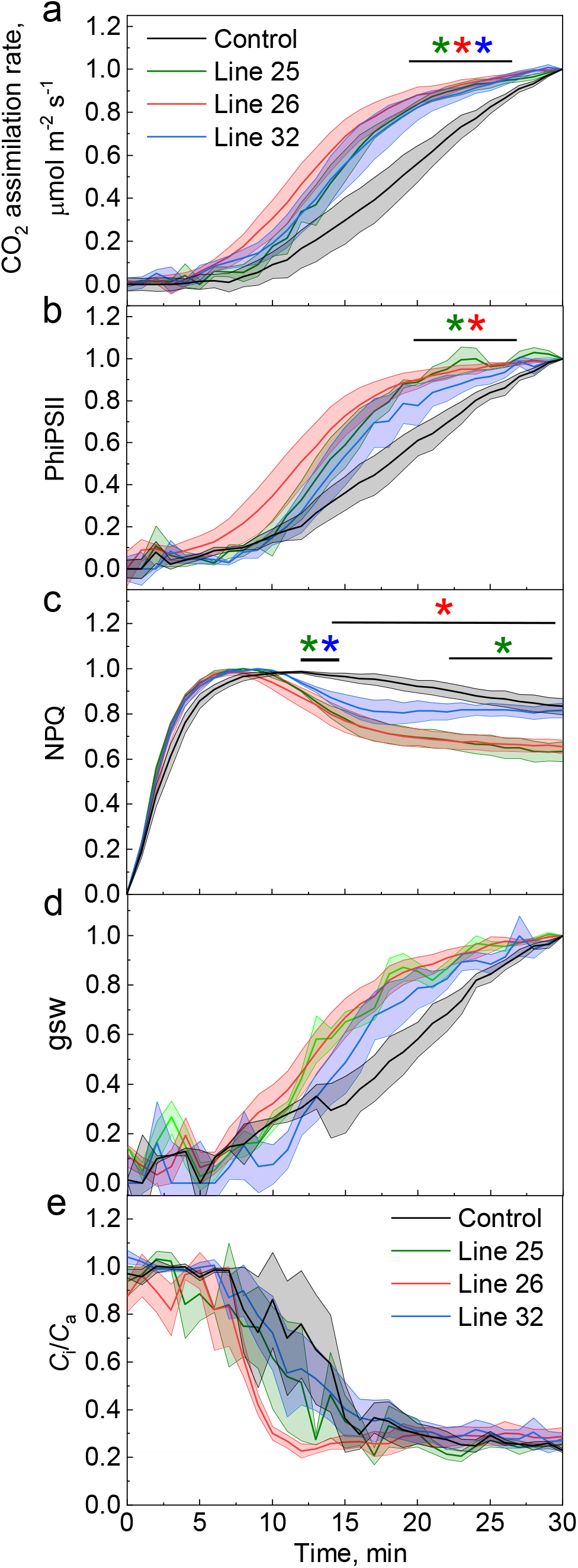
Induction of photosynthesis during the first 30 min of illumination with actinic light of 1000 µmol m^-2^ s^-1^ in control plants and transgenic sorghum lines overexpressing Rieske. PhiPSII, the effective quantum yield of PSII; NPQ, non-photochemical quenching; gsw, stomatal conductance to water vapor; *C*_i_/*C*_a_, the ratio between the intercellular and ambient CO_2_ partial pressures. The values of each parameter, except for *C*_i_/*C*_a_, at the beginning and end of the 30-min illumination were normalised to 0 and 1, respectively, to facilitate comparison of the kinetics. Azygous plants were used as control. Mean ± SE, *n* = 5 biological replicates. Asterisks indicate intervals of statistically significant differences between transgenic and control plants (one-way ANOVA and Dunnett’s post-hoc test at *P* < 0.05).

## Discussion

Sorghum is one of the most important crops in the world which serves as a source of food, fodder and fuel. Sorghum can withstand severe droughts allowing it to grow in regions where other major crops cannot be grown, like Sub-Saharan Africa. However, in recent years, genetic progress in sorghum yield has stagnated and not kept pace with increasing demand (Ananda et al., 2020). Therefore, it is critical to develop new approaches for increasing sorghum productivity. Based on crop model predictions, up to 10% improvement in sorghum yield could be harnessed from improving photosynthesis (Wu et al., 2019). According to the biochemical model of C_4_ photosynthesis, assimilation at low *p*CO_2_ is limited by PEPC and CA activities and mesophyll conductance to CO_2_, whilst assimilation at ambient and high *p*CO_2_ is limited by Rubisco, electron transport or the regeneration rate of Rubisco’s substrate (von Caemmerer, 2000; von Caemmerer, 2021; von Caemmerer and Furbank, 1999). Contribution of some of these factors to C_4_ photosynthesis and plant productivity was recently tested using transgenic approach in the model C_4_ plant *S. viridis* (Alonso-Cantabrana et al., 2018; Ermakova et al., 2022; Osborn et al., 2016), and improvements of C_4_ photosynthesis were shown in *S. viridis* and *Z. mays* with increased Rieske and Rubisco content (Ermakova et al., 2019; Ermakova et al., 2021b; Salesse-Smith et al., 2018). Here we expand on our previous results and assess how Rieske overexpression affects productivity of sorghum.

Rieske overexpression in sorghum provided increased Cyt*b*_6_*f* activity, confirmed by monitoring the build-up of NPQ and dynamics of electron transport during the photosynthesis induction. A faster build-up of NPQ during dark-light transition in transgenic plants indicated larger ΔpH due to increased Cyt*b*_6_*f* activity (Fig. 5c). Observing this transient increase is possible because NPQ engages on the scale of seconds to minutes (Müller et al., 2001). However, by the end of 3-min illumination periods, *pmf*, which is a sum of ΔpH and Δψ (the membrane potential), did not differ between genotypes (Fig. 5a). This was consistent with the largely unchanged electron transport parameters and CO_2_ assimilation rates (Fig. 3 and Fig. 4) – all indicating that steady-state rates of electron transport were unaltered in sorghum plants overexpressing Rieske. Similar observations were made with tobacco plants overexpressing Rieske (Heyno et al., 2022). In C_3_ plants, this phenomenon could be explained by a conserved relationship between ΔpH and NPQ which would reduce PSII activity in case if electron transport rate exceeds the capacity of dark reactions of photosynthesis to consume ATP and NADPH, typically due to a limited availability of CO_2_ (Kanazawa and Kramer, 2002). C_4_ photosynthesis, however, is less limited by CO_2_ due to the C_4_ cycle concentrating CO_2_ around Rubisco, and an increase of electron transport rate is projected to provide proportional increase in assimilation (von Caemmerer and Furbank, 2016). Increasing Rieske content in *S. viridis* was sufficient to enhance electron transport rates, resulting in higher photosynthesis at non-limiting CO_2_ and high light (Ermakova et al., 2019). One possible explanation for the difference observed between *S. viridis* and sorghum overexpressing Rieske is that C_4_ crops underwent a selection for photosynthetic traits which could have altered a balance between electron transport components. For example, translational efficiency of Lhca6, a subunit of the light-harvesting complex I that facilitates a formation of PSI supercomplex involved in cyclic electron flow (Otani et al., 2018), was significantly enhanced during maize domestication (Zhu et al., 2021). Changes to the ratio of cyclic to linear electron flow likely resulted in additional or different factors limiting electron transport. Better understanding of photosynthetic changes that C_4_ crops underwent during domestication will help to uncover additional targets for accelerating steady-state electron transport and photosynthesis rates (Hu et al., 2018; Tao et al., 2020a; Zhu et al., 2021). Increased assimilation rates detected in maize with increased Rubisco content at non-limiting CO_2_ and high light could also be indicative of an altered relationship between electron transport and Rubisco limitations in C_4_ crops (Salesse-Smith et al., 2018).

Interestingly, the largest difference in electron transport and assimilation between plants overexpressing Rieske and control plants was detected during the induction of photosynthesis. Induction, or activation of photosynthesis during dark-to-light transition, is a complex process that, in C_3_ plants, requires opening of stomata, activation of Rubisco and other enzymes and a build-up of metabolite concentrations (Deans et al., 2019; Slattery et al., 2018). Faster activation of photosynthesis was identified as one of the desirable traits in crop plants that could allow up to 20% increase in total assimilation (Acevedo-Siaca et al., 2020; Long et al., 2022). Activation of C_4_ photosynthesis is further complicated by the distribution of electron transport and metabolic reactions between mesophyll and BS cells and a necessity to coordinate activities of C_4_ and C_3_ cycles (Furbank and Taylor, 1995; Kromdijk et al., 2014). Our results support previous works suggesting that, due to the operation of carbon concentrating mechanism, activation of C_4_ photosynthesis is less limited by stomatal conductance compared to C_3_ photosynthesis (Furbank and Walker, 1985; Usuda and Edwards, 1984). Indeed, in our experiments, *C*_i_/*C*_a_ never dropped below 0.25 (Fig. 6e) which is equivalent to *C*_i_ ≥ 100 µmol mol^-1^ sufficient to saturate assimilation (Pignon and Long, 2020). Instead, the observed increase of CO_2_ assimilation rates during the light-induced activation of photosynthesis in sorghum overexpressing Rieske was underpinned by the higher PhiPSII (Fig. 6). It is conceivable that activation of photosynthetic enzymes provides a strong sink for ATP and NADPH, and an increased Cyt*b*_6_*f* activity could transiently stimulate electron transport to activate enzymes and build-up metabolite gradients faster. In line with this, activation of Rubisco that uses ATP was suggested as one of the major factors limiting C_4_ photosynthesis during the induction (Wang et al., 2021).

Sorghum plants with increased Rieske content had increased biomass and grain yield. Transgenic plants produced more leaves and had larger leaves during the vegetative growth phase and accumulated more biomass by the end of growth season, compared to control plants (Fig. 2, Fig. S2). Moreover, transgenic plants produced about 20% more grain by weight and number (Fig. 2g and Fig. 2h) indicating that higher yield of transgenic plants was largely attributed to setting more seeds. Grain yield and grain number highly correlate in sorghum, and a supply of assimilates during the seed setting largely determines grain number (Craufurd and Peacock, 1993; van Oosterom and Hammer, 2008). Although the conditions of induction experiment (Fig. 6) were designed to maximise differences between control and transgenic plants and did not occur in glasshouse conditions, a cumulative effect of transiently or marginally increased electron transport rates could explain the observed improvements in biomass and yield of transgenic plants. Similarly, small changes in CO_2_ assimilation rates due to accelerated NPQ relaxation were shown to result in significant increase in biomass of tobacco grown in the field (Kromdijk et al., 2016). Taken together, our results show that increasing Rieske content in sorghum improves the light use efficiency and stimulates biomass production therefore presenting a promising trait to be introduced into commercial sorghum varieties. Exploring variation of Rieske content in sorghum variety panels (Tao et al., 2020b) could inform breeding programs for developing higher yielding sorghum with improved photosynthesis.

## Conclusion

In sorghum, effects of Rieske overexpression on stimulating electron transport were more apparent during transient photosynthetic responses suggesting that other photosynthetic components co-limit the steady-state electron transport rate. However, faster responses of electron transport and CO_2_ assimilation to light transients resulted in plants overexpressing Rieske accumulating more biomass and producing more grain. Our results indicate that increasing Rieske content, relative to other electron transport components, is a promising way to boost productivity of C_4_ crops and ensure food and energy security.

## Materials and Methods

### Generation and selection of transgenic plants

The gene construct for Rieske overexpression was created using the Golden Gate cloning system (Engler et al., 2014), as described in Ermakova et al. (2019). The first expression module containing the selection marker was occupied by *neomycin phosphotransferase II* (*ntpII*) driven by the *Z. mays ubiquitine1* promotor. The second expression module contained the coding sequence of *petC* from *Brachipodium dystachion* (*BdpetC*) under the control of the *Z. mays ubiquitine1* promotor. The construct was transformed into *Agrobacterium tumefaciens* strain AGL1 and then into *Sorghum bicolor* Tx430 according to (Gurel et al., 2009). Transgenic plants were analysed for the *ntpII* copy number by digital PCR (iDNA genetics, Norwich, UK). Azygous plants were used as control in all experiments. Transcript abundance of *BdpetC* was estimated by qPCR as described in Ermakova et al. (2019). Electron transport at ambient light intensity and leaf properties (relative chlorophyll and leaf thickness) were assayed with MultispeQ (Kuhlgert et al., 2016) and analysed using the PhotosynQ platform (https://photosynq.com).

### Plant growth conditions

Plants were grown in a glasshouse at ambient irradiance, 28 °C day, 20 °C night and 60% humidity. Plants were shuffled every 2-3 days to reduce any positional growth effects. The youngest fully expanded leaves of the 4-5 weeks old plants were used for all physiological analyses. Photos of plants, leaves and tiller count and height measurements were done during the fully emerged leaves stage, 5 weeks after germination.

### Gas exchange and fluorescence analyses

Rates of CO_2_ assimilation were measured over a range of *p*CO_2_ and irradiance using a portable gas-exchange system LI-6800 (LI-COR Biosciences, Lincoln, NE). Chlorophyll fluorescence was assessed simultaneously with a Fluorometer head 6800-01 A (LI-COR Biosciences). Leaves were first equilibrated at 400 ppm CO_2_ in the reference side, 1500 µmol m^−2^ s^−1^, leaf temperature 28 °C, 60% humidity and flow rate 500 µmol s^−1^. CO_2_ response curves were conducted under the constant irradiance of 1500 µmol m^−2^ s^−1^ by imposing a stepwise increase of *p*CO_2_ from 0 to 1600 ppm at 3 min intervals. Light response curves were measured at the constant *p*CO_2_ of 400 ppm in the reference cell under a stepwise decrease of irradiance from 0 to 2000 µmol m^−2^ s^−1^ at 3 min intervals. Red-blue actinic light (90%/10%) was used in all measurements. The effective yield of PSII (PhiPSII) was assessed at the end of each step upon the application of a multiphase saturating pulse (8000 µmol m^−2^ s^−1^) and calculated according to Genty et al. (1989).

Induction of photosynthesis was analysed on dark-adapted overnight plants. First, leaves were clipped into LI-6800 chamber in darkness and the minimum and maximum levels of fluorescence were recorded upon the application of a saturating pulse. After that, leaves were illuminated with actinic light of 1000 µmol m^−2^ s^−1^, and gas-exchange and fluorescence parameters were recorded every 1 min for 30 min. NPQ was calculated according to Bilger and Björkman (1990). All parameters were normalised for min and max values to facilitate comparison of the kinetics.

### Electron and proton transport

Fluorescence parameters informing on the activity of PSII and absorbance at 820 nm reflecting the formation of oxidised P700, the reaction centre of PSI, were analysed simultaneously by the Dual-PAM-100 (Heinz Walz, Effeltrich, Germany). Measurements were done using red actinic light and 300-ms saturating pulses of 10000 µmol m^−2^ s^−1^. Leaves were first dark-adapted for 30 min to record the minimal and maximal levels of fluorescence and calculate F_V_/F_M_, the maximum quantum yield of PSII. Then, a saturating pulse was applied after a pre-illumination with strong far-red light to record the maximal level of P700^+^ signal and, after the pulse, the minimal level of P700^+^ signal. Next, leaves were illuminated for 10 min with an actinic light of 378 µmol m^−2^ s^−1^. After that, photosynthetic parameters were assessed over a range of irradiances from 0 to 2043 µmol m^−2^ s^−1^ at 2 min intervals by applying a saturating pulse at the end of each step. The effective quantum yield of PSII (PhiPSII), the yield of non-photochemical quenching (PhiNPQ) and the yield of non-regulated non-photochemical reactions within PSII (PhiNO) were calculated according to Kramer et al. (2004). The effective quantum yield of PSI (PhiPSI), the non-photochemical yield of PSI caused by the donor side limitation (PhiND) and the non-photochemical yield of PSI caused by the acceptor side limitation (PhiNA) were calculated according to Klughammer and Schreiber (2008).

The electrochromic shift signal (ECS) was monitored as the absorbance change at 515-550 nm using the Dual-PAM-100 equipped with the P515/535 emitter-detector module (Heinz Walz). The absorbance signal at 535 nm was monitored simultaneously. Leaves were first dark-adapted for 40 min and the amplitude of ECS induced by a single turnover flash was recorded. Dark-interval relaxation kinetics of ECS was then recorded after 3-min intervals of illumination with red actinic light of increasing irradiance. Proton motive force (*pmf*) was estimated from the amplitude of the rapid decay of ECS signal upon light-dark transition, normalised for the ECS induced by the single turnover flash. Proton conductivity of the thylakoid membrane through ATP synthase was calculated as an inverse time constant obtained by the fitting of first-order ECS relaxation kinetics after Sacksteder and Kramer (2000).

### Protein isolation and Western blotting

Total protein extracts were isolated from 0.7 cm^2^ frozen leaf discs as described in Ermakova et al. (2019) and separated by SDS-PAGE. Proteins were then transferred to a nitrocellulose membrane and probed with antibodies against various photosynthetic proteins in dilutions recommended by the producer: Rieske (AS08 330, Agrisera, Vännäs, Sweden), D1 (Agrisera, AS10 704), PGR5 (Agrisera, AS163985). Quantification of immunoblots was performed with Image Lab software (Biorad, Hercules, CA).

### Statistical analysis

The relationship between mean values of different groups was tested by one-way ANOVAs with Dunnett’s or Tukey’s *post-hoc* test or by two-tailed heteroscedastic *t*-test, as indicated in figure legends.

## Acknowledgements

We thank Spencer Whitney for the gift of RbcL antibody and Siena Mitchell, Alexandra Williams, Ayla Manwaring, Kelly Chapman and James Samuel Nix for technical assistance. This work was performed under the collaborative project agreement between the Australian National University and CSIRO Food & Agriculture and supported by the Australian Research Council Centre of Excellence in Translational Photosynthesis (CE140100015) and by the seed grant from the Research School of Biology at the Australian National University. Authors declare no conflict of interests.

## Author Contributions

SvC, ME and RTF designed the research; SB generated transgenic plants; ME, RW, SB and ZT performed research; ME and RW analyzed data; ME wrote the paper.

## Short legends for Supporting Information

### Supplementary figures

**Fig. S1.**
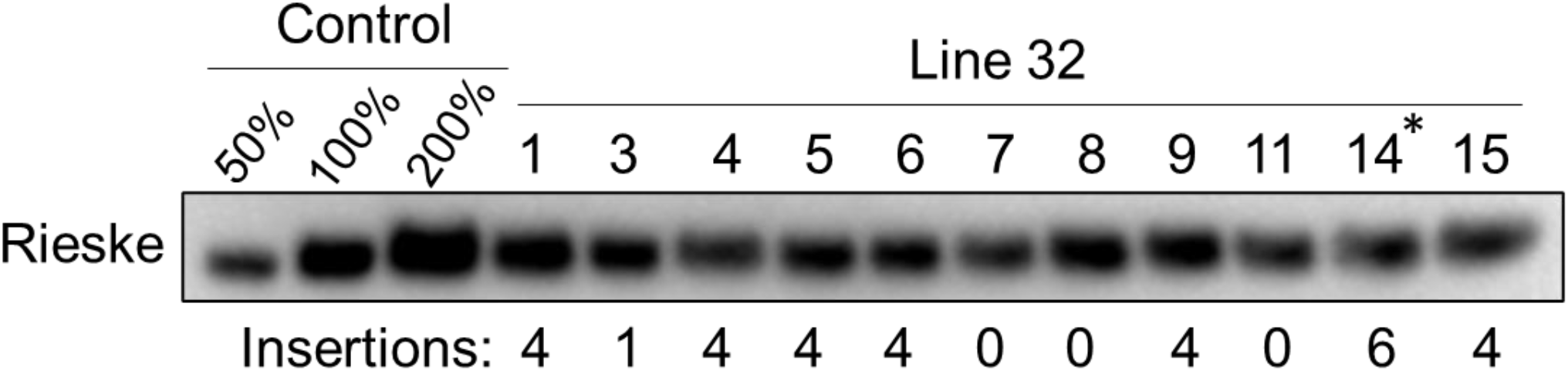
Immunodetection of Rieske in the T_1_ progeny of line 32. Samples were loaded on leaf area basis, and the WT sample was used as a control. Insertion numbers indicate a copy number of *ntpII* obtained by digital PCR. Asterisk indicates homozygous plant 32-14 which T_2_ progeny was used in further experiments.

**Fig. S2.**
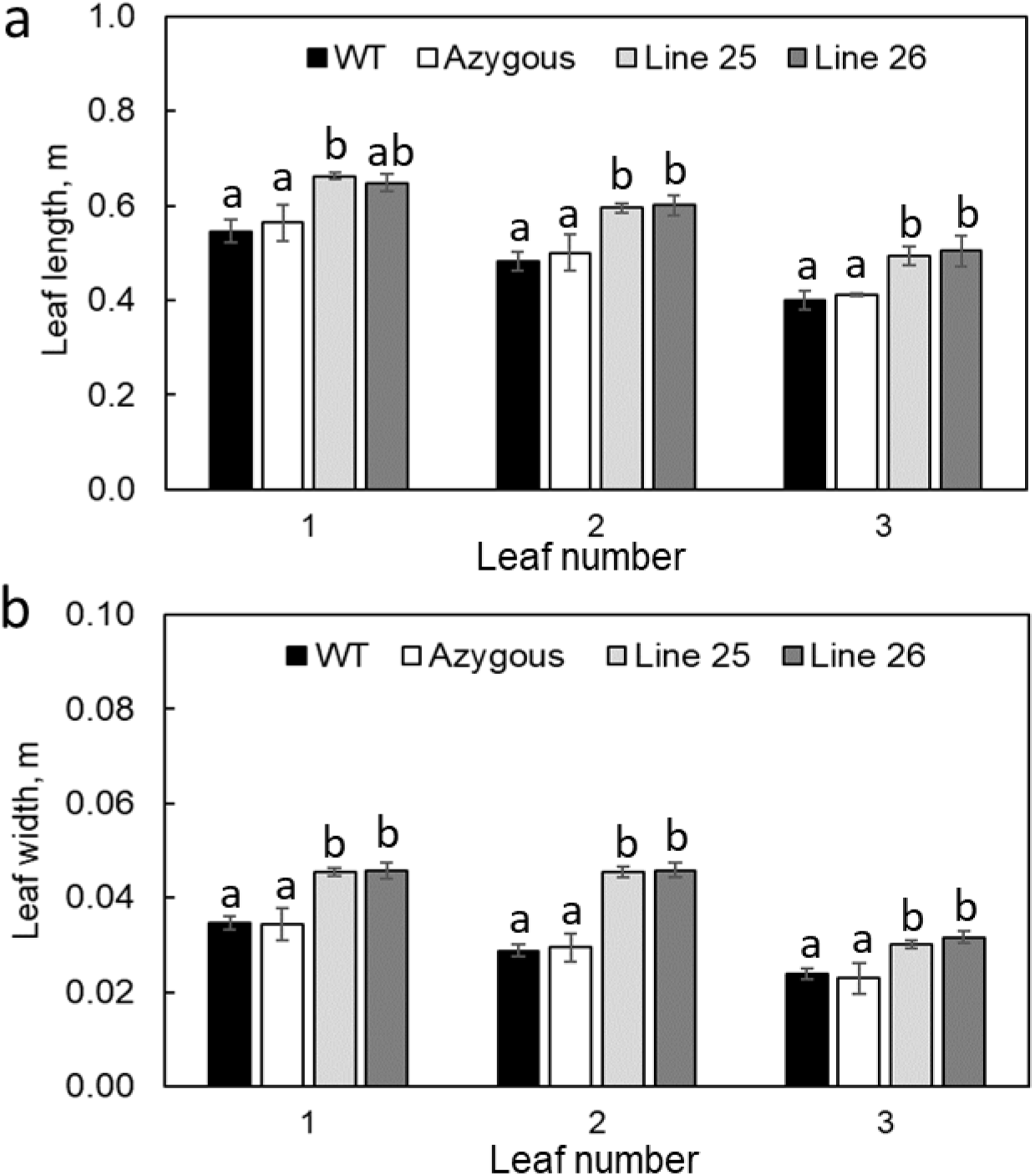
Length and width of top fully expanded leaves from WT, azygous and Rieske overexpressing plants. Mean ± SE, *n* = 5 biological replicates. Letters indicate statistically significant differences between groups (one-way ANOVA and Tukey post-hoc test, *P* < 0.05).

